# Detecting global influence of transcription factor interactions on gene expression in lymphoblastoid cells using neural network models

**DOI:** 10.1101/2021.06.18.449031

**Authors:** Neel Patel, William S. Bush

## Abstract

**Background:** Transcription factor(TF) interactions are known to regulate target gene(TG) expression in eukaryotes via TF regulatory modules(TRMs). Such interactions can be formed due to co-localizing TFs binding proximally to each other in the DNA sequence or over long distances between distally binding TFs via chromatin looping. While the former type of interaction has been characterized extensively, long distance TF interactions are still largely understudied. Furthermore, most prior approaches have focused on characterizing physical TF interactions without accounting for their effects on TG expression regulation. Understanding TRM based TG expression regulation could aid in understanding diseases caused by disruptions to these mechanisms. In this paper, we present a novel neural network based TRM detection approach that consists of using multi-omics TF based regulatory mechanism information to generate features for building non-linear multilayer perceptron TG expression prediction models in the GM12878 immortalized lymphoblastoid cells.

**Results:** We estimated main effects of 149 individual TFs and interaction effects of 48 distinct combinations of TFs forming TRMs based on their influence on TG expression. We identified several well-known and discovered multiple previously uncharacterized TF interactions within our detected set of TRMs. We further characterized the pairwise TRMs using long distance chromatin looping and motif co-occurrence data. We found that nearly all the TFs constituting TRMs detected by our approach interacted via chromatin looping, and that these TFs further interacted with promoters to influence TG expression through one of four possible regulatory configurations.

**Conclusion:** Here, we have provided a framework for detecting TRMs using neural network models containing multi-omics TF based regulatory features. We have also described these TRMs based on their regulatory potential along with presenting evidence for the possibility of TF interactions forming the TRMs occurring via chromatin looping.

## Background

Concerted and combinatorial binding of transcription factors (TF) within the *cis*-regulatory elements of target genes (TG) in humans gives rise to transcriptional regulatory modules(TRMs), which are essential for regulating TG expression[1]. These TRMs influence TG expression both additively and non-additively as seen in model systems[2], [3]. Physical interaction among TFs, which is the basis for the formation of these TRMs, has been theorized to occur based on different models[4]. “Enhanceosome” and “Billboard” models represent linear co-operative interactions among TFs brought about by their DNA sequence based motif proximity, with “Billboard” models allowing a more flexible motif orientation and spacing than the “Enhanceosome” [4][5]. Alternatively, the “TF collective” model comprises of non-linear TF interactions, independent of the DNA sequence motif composition. While this model was originally based in part on protein-protein interactions among TFs, it may also be due to distally binding TFs brought together via chromatin looping[4]. Previous computational approaches have mainly focused on characterizing TF interactions using the “Enhanceosome” and “Billboard” models[6]–[9]. However, the influence of TFs interacting via the non-linear “TF collective” model on TG expression is not well understood. Disruption in TRM based TG expression regulation, caused by genetic mutations in the TFs forming these TRMs, has been associated with several diseases. For instance, genes encoding TFs forming the BAF chromatin remodeling complex and the cohesin complex are found mutated in some congenital disorders[10]. Similarly, mutations in the TFs mediating the interaction of distally binding TFs with TG promoters and in those present within the heterochromatin forming polycomb-repressive complex(PRC) have been shown to cause different types of tumors[11]. Genetic disruption in the AP-1 factor complex based TG regulation has been found to cause neurodevelopmental disorders as well as autoimmune diseases [12][13]. While most of these examples have been studied in isolation, a systems-wide understanding of TG expression regulation driven by TRMs will likely unravel regulatory mechanisms underlying a range of other diseases.

Availability of high-throughput ChIP-Seq datasets, which provide the sequence specific binding information for each TF, has enabled researchers to detect TF interactions across the whole genome. Gerstein *et al*. analyzed the co-localization maps of different TFs in K562 and GM12878 cell lines to detect significantly co-associating TFs using a discriminative machine learning approach[6]. They detected several well-characterized TF interactions such as the GATA1-complex(GATA1-GATA2-TAL1), MYC complex(MYC-MAX-E2F6) and the AP1-factors (FOS-JUN-JUND-FOSL) as well as some novel TF interactions such as GATA1-CCNT2–HMGN3 and GATA1-NRSF-REST using their approach. Others have used non-parametric modeling approaches to identify pairwise or higher-order interactions of TFs[7], [9]. For example, Guo and Gifford developed a topic modeling approach called Regulatory Motif Discovery(RMD) that identifies different TF interactions utilizing TF co-localization information.[7] They detected multiple well known TF interactions such as the cohesin complex (CTCF-RAD21-SMC3) complex, the transcription pre-initiation complex (POL2-TBP-TAF1) and the AP1 factor complex. Bailey *et. al*. identified several literature-annotated interactions by identifying closely binding TFs based on significant spacings between their sequence motifs[8]. Lastly, soft and hard clustering methods such as k-means clustering, non-negative matrix factorization and self-organizing maps have also been used to identify co-localizing TFs across the genome[14]–[17]. Although these studies have helped in systems-wide detection and characterization of TF interactions, they have the following limitations: 1) Interactions that non-additively influence TG expression via distally binding TFs caused by chromatin looping (the “TF collective” model) cannot be detected using the abovementioned methods, as they rely upon TF co-localization information to identify proximally co-binding TFs (more consistent with the “Enhanceosome” and “Billboard” models). 2) The unsupervised clustering and topic modelling methods require the user to pre-determine the number of TF interactions to be identified preventing the agnostic discovery of TF interactions. 3) Lastly, and most importantly, for most of these studies the quantitative impact of TF interactions on TG expression remains unknown.

In this study, we use a multi-omics machine learning framework to model the impact of multiple TF based regulatory mechanisms on TG expression and detect TRMs based on the interaction effects learned from these models. We generated a gene regulatory network(GRN) containing information from datasets representing TF-TG, TF cooperativity and TG co-regulation. The TF-TG interactions in our multi-omic GRN were also weighted based on chromatin looping interactions made by distally binding TFs with the TG promoters to appropriately capture their effect on TG regulation. We used the features from this GRN to predict TG expression values in the GM12878 lymphoblastoid cell line(LCL) using non-linear deep learning multilayer perceptron(MLP) prediction models. By aggregating interaction effects among different combinations of TFs from our learned models, we were able to identify specific TRMs that had high impact on TG expression. We validated the TF interactions, that we discovered within these TRMs, based on long distance chromatin looping contacts between their distal binding sites and significant spacing between their motifs for proximal binding sites. We also characterized the transcriptional regulatory programs for these modules based on the orientation and interaction of the corresponding ChIP-seq peaks relative to the promoters of TGs. Using our flexible multi-omics machine learning framework, we were able to detect TRMs significantly influencing TG expression, while characterizing their regulatory architectures using biologically relevant information.

## Results

### Target gene expression could be better predicted by modelling complex non-linear interactions among transcription factors

We hypothesized that information beyond sequence co-localization of TFs would be useful for detecting TRMs, formed by the “TF collective” model described in **Background**, essential for TG expression regulation. As a basis to examine this hypothesis, we developed a multi-omics machine learning framework shown in ***Figure-1***, which modelled the influence of multiple TF based regulatory mechanisms (TF co-operativity, TF-TG binding and TF-TG co-regulation) on TG expression, in the GM12878 LCL, by using features derived from a gene regulatory network(GRN) built using the PANDA algorithm[18]. We have shown previously that features obtained from such a GRN explain more variance in TG expression compared to using TF-TG binding information alone.[19]

**Figure 1:**
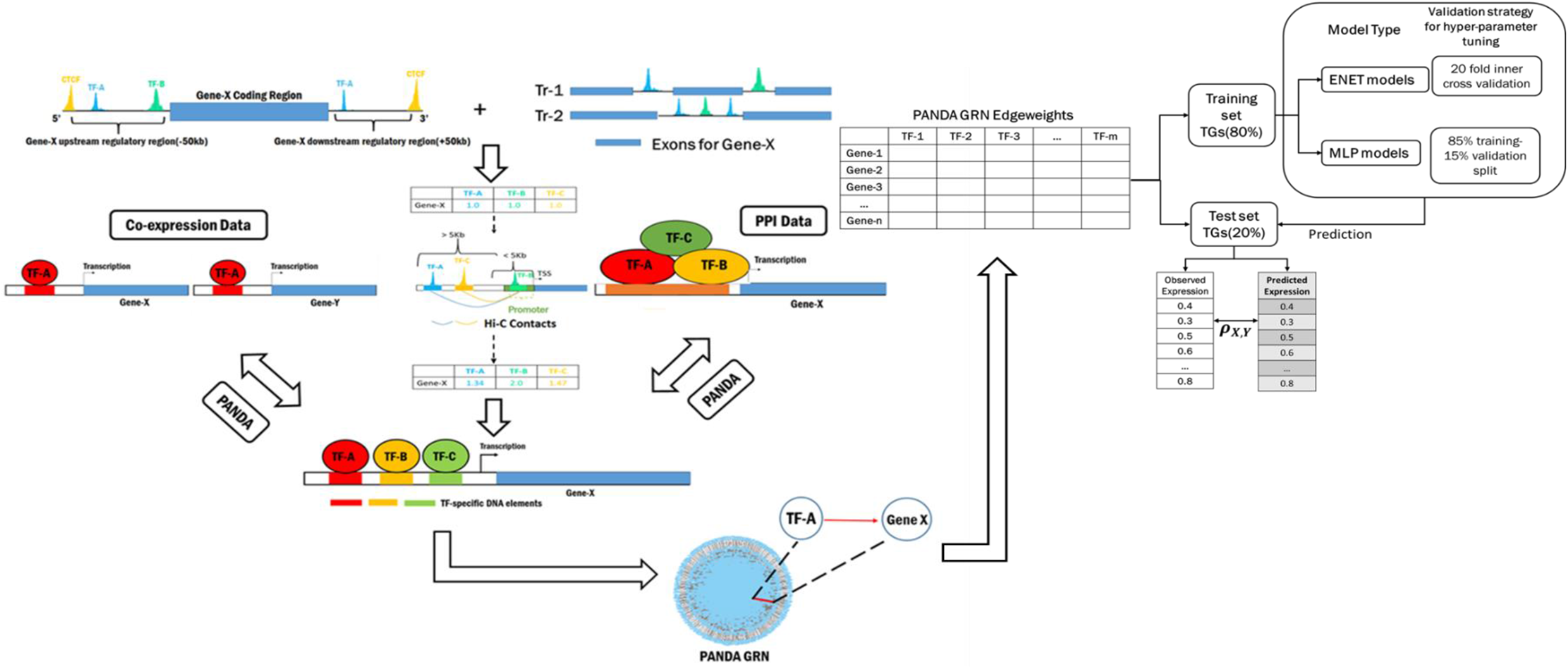
Using a multi-omics GRN framework to predict gene expression. We downloaded ChIP-seq data for 149 GM12878 TFs from the ENCODE consortium whose accession numbers are provided in Supplementary table S1. We used the peaks that passed the optimal IDR(Irreproducible Discovery Rate) threshold defined by the consortium and mapped them onto the regulatory region of each gene to define TFBs. We used CTCF peaks within a 50Kb window upstream and downstream of the gene body in order to demarcate the regulatory boundaries. Furthermore, we weighted the TF-TG interactions based on the number of contacts made by the corresponding peaks with the promoter of TGs. We used a weighting scheme where promoter TFBS were automatically up-weighted because of the inability of HiC data to capture them due to limited resolution. We created PANDA GRNs using the weighted adjacency matrices, the PPI data corresponding to the TFs obtained from BioGRID and the lymphoblastoid co-expression data obtained from GEUVADIS. After generating the PANDA GRN, we built elastic net(ENET) and multilayer perceptron(MLP) models that used them as input features to predict log FPKM values(gene expression) of an independent dataset. We used two different internal cross-validation strategies to train the ENET and the MLP models and assessed their accuracy by computing Pearsons correlation coeffcient(PCC) between observed and predicted expression.

In order to generate these GRNs, we first extracted all the *cis*-regulatory and intronic TF binding sites(TFBS) corresponding for 149 TFs for each TG. Next, we created an adjacency matrix weighted by the number of chromatin looping interactions between each TF and the TG promoter based on GM12878 high throughput chromatin capture(Hi-C) data. We used this weighted adjacency matrix to create a motif network reflecting GM12878 specific co-expression and PPI networks.

We used the TF-TG edge-weights from this GRN to build our prediction models where we tested for purely additive as well non-additive influence of TF features on TG expression. As a baseline comparison, we first built ElasticNet(ENET) regularized regression models which assume an additive(linear) influence of TF features over TG expression without considering any non-additive effects of TF interactions. We next used a traditional neural network based multilayer perceptron(MLPs) capable of modelling non-additive(non-linear) interaction effects of TFs on TG expression, which we hypothesized will help identify “TF collective” based TRMs. We further evaluated a hybrid model (MLP-U) that can decompose the effects into additive and non-additive components. This model was composed of set of univariate MLP models capturing individual TF influence over TG expression along with a traditional MLP to capture all possible interaction effects (see ***Supplementary Figure S1***). Thus, we used 3 different prediction models:1) an ENET to model TF main effects only 2) an MLP to model complex interaction effects, and 3) an MLP-U model that can be decomposed into additive and nonadditive components. Further details about these models, especially the MLP and MLP-U architectures have been provided in the **MLP network architecture and building the prediction models** section of the **Methods**.

We used an independent GM12878 LCL expression dataset(accession: ENCSR889TRN) to train and test the prediction models. We used 80% of the total TG set for training the models as well as for internal cross validation and used the remaining 20% for testing the prediction accuracy (***Figure 1***). We performed the prediction task for 20 iterations, each time using a different set of test TGs. We used the median Pearson’s correlation coefficient(PCC) aggregated across all of these iterations to compare the performance of the 3 different models. As shown in ***Figure-2A***, the models capable of capturing interaction effects of TFs (MLP and MLP-U) perform significantly better (median PCC_MLP_ = 0.68; median PCC_MLP-U_ = 0.63) compared to the ENET models (median PCC_ENET_ = 0.57), which model linear influence of individual TFs on gene expression. This improvement in performance was also statistically significant (median PCCMLP vs. median PCC_ENET_ p-value = 1.91e-06; median PCC_MLP-U_ vs. median PCC_ENET_ p-value = 1.91e-06) as calculated by performing paired Wilcoxon sign rank tests. We have shown the PCC for the three types of models obtained from each prediction iteration in ***Supplementary table S2*.** Furthermore, we observed that over all the prediction iterations for the MLP-U models, the main effects, obtained from their univariate component, were more predictive of TG expression, explaining about 34% of the variance on average, than the interaction effects, captured by their MLP component, which explained 23% of the variance in TG expression(see **Partitioning TG expression variance explained by the univariate and MLP components from the MLP-U models** of ***Supplementary Methods*** and ***Supplementary Figure S2***).

**Figure 2:**
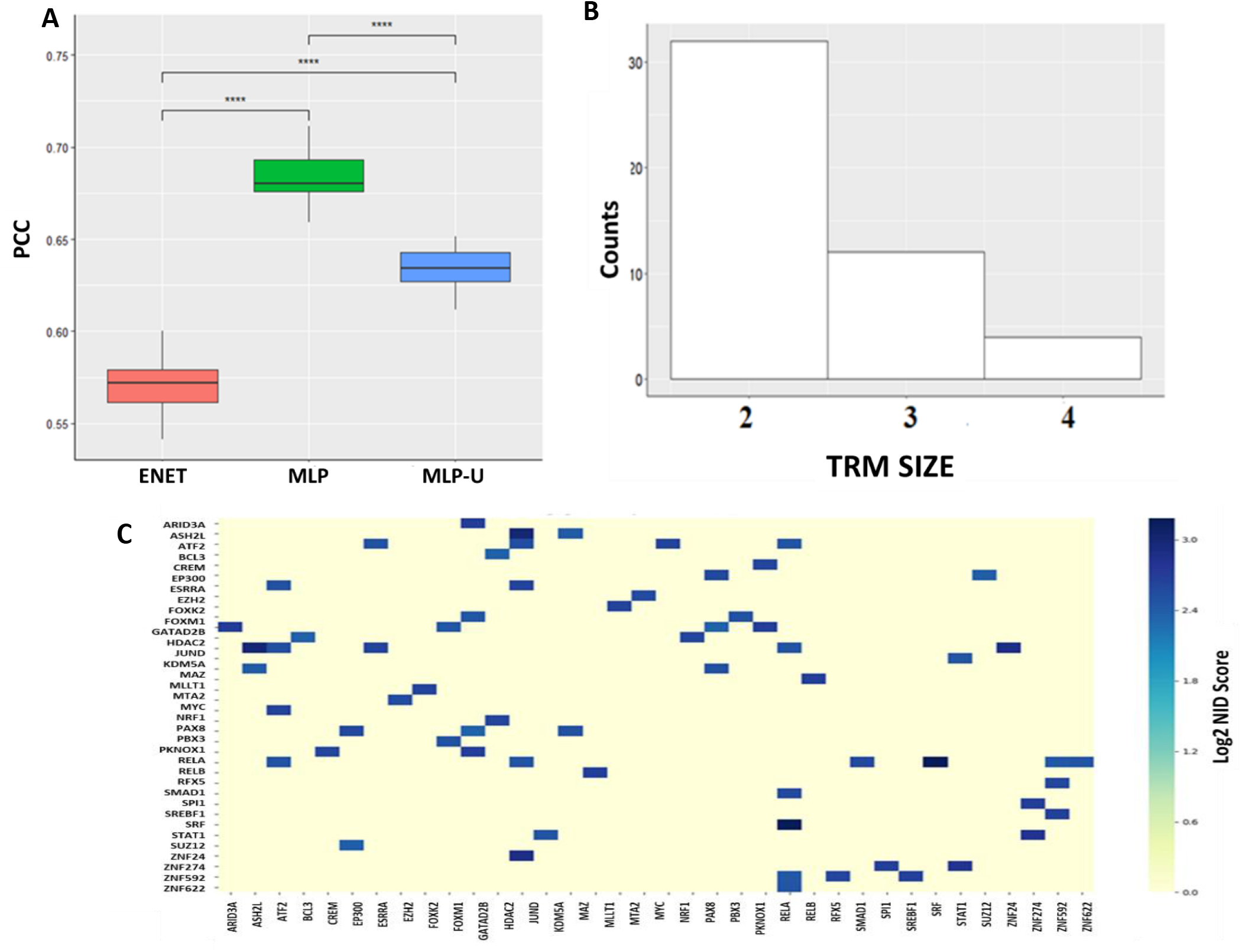
Learning global transcriptional regulatory patterns from multi-omics GRN based machine learning approach. A) Boxplot showing the performance of the MLP, MLP-U and ENET models obtained from the prediction of 2356 TGs over 20 iterations (****p-value < 0.0001). B) Barplot showing the sizes of the 48 TRMs, defined as the number of TFs in each of them, detected from the learned MLP-U models based on the NID algorithm. C) Heasumap showing the strength of the interactions for 32 pairwise TFs calculated based on the Log2 NID scores.

In conclusion, accurate prediction of TG expression requires efficient modelling of main effects of individual TFs as well as interaction effects of TRMs.

### Context dependent influence of individual transcription factors on target gene expression could be discerned from our models

The MLP-U architecture, described in ***Supplementary Figure S1***, allowed us to model main effects of individual TFs separately from the interaction effects of TF combinations. We used equations(1)–(18) to calculate these main and interaction effects from the trained MLP-U models(see **Obtaining main and interaction effects from the MLP-U models** of the **Methods** section).

Our learned MLP-U models contained individual univariate MLPs corresponding to each one of the 149 TFs. We aggregated all the learned connection weights at the first layer of these MLPs and multiplied them with the nodal influence score for each node in that layer. After averaging these nodal scores, we calculated an average main effect for each TF across all the prediction iterations followed by scaling it in the range (−1,1). We have provided the scaled and the raw main effects for each of the 149 TFs **(*Supplementary table S3A***).

In order to examine the validity of our main effects aggregation approach, we divided the TFs into 5 bins based on their scaled main effects (***Supplementary Figure S3*** and ***Supplementary table S3A***). The bin placement of the TFs derived from their main effects reflected their functional roles. For instance, activating TFs such as TAF1, MYC, TBLXR1, RELA and BCL11A were present in the right most bin(5) because of their highly positive main effects. On the other hand, transcription repressors such as MXI1, HDAC2, SMC3, MAZ and ZNF592 had strongly negative main effects placing them in the left most bin (1). We compared the main effects obtained from the MLP-U models to those obtained from the ENET model by computing the difference in ranks(DIR) of the TFs based on their effects for the two modelling approaches (***Supplementary table S3A***). Positive DIR for a TF reflected decrease in the MLP-U main effect, while a negative DIR represented increase in the MLP-U main effect, compared to that obtained from the ENET models.

We found that TFs with extremely negative DIR based on their MLP-U main effects, such as ZNF143(−128), TBLXR1(−121), DPF2(−115), E4F1(−115) and YY1(−110), were transcriptional activators in specific contexts representing their interactions with other TFs[20]–[22]. Alternatively, TFs with extremely positive DIR, such as ZBTB40(112), HDAC2(112), SIN3A(124), SMAD1(98) and KDM1A(125) could act as repressors when interacting with other TFs[23]–[27]. We also found an extremely positive DIR for the well-known transcriptional activator TBP(125), which requires other promoter binding TFs such as the TBP-associating factors(TAFs) to recruit RNA polymerase II and to exert its effect[28]. Thus, while ENET models captured influence of TFs assuming independent effects on TG expression, main effects obtained from the MLP-U models are adjusted for context in which the TF binding event occurs (see equation (1)).

### Interaction effects aided the detection of well-known and novel transcription factor regulatory modules

The MLP component of the MLP-U models quantify the non-additive interaction effects of different combinations of TFs on TG expression. These effects could reflect the influence of non-linear “TF collective” interactions on TG expression. We applied the NID algorithm[29] to compute interaction effects in the form of NID scores for such TRMs. This calculation is done at each node of the first layer, for all the possible combinations/orders of the interactions and only the top ranked interactions for each order are retained. The interactions are aggregated such that lower order redundant interactions are removed and higher order top ranking interactions are retained giving a final set of highly impactful interactions of different orders. We defined these interactions along with their average NID scores as TRMs. We applied Log2 normalization to the average NID scores calculated for each TRM across all the 20 prediction iterations (***Supplementary table S3B***).

We detected 48 unique TRMs out of which 32 were pairwise interactions, 12 were 3-way and 4 were 4-way interactions as shown in ***Figure-2B***. The pairwise TRMs were formed by 36 unique TFs, 3-way TRMs were formed by 22 TFs and the 4-way TRMs were formed by 12 TFs. Furthermore, we observed that among the higher order (3-way or higher) TRMs, the “nestedness” or the proportion of all the possible pairwise TRMs being also detected was never 100%(***Supplementary Figure S4***). We found that JUND formed the largest number of TRMs(11) followed by GATAD2B(10), RELB(10) and ATF2(9). All of these TFs are versatile DNA binding proteins capable of affecting cell proliferation, division and apoptosis, which explains their presence in a large number of TRMs.

Multiple literature annotated TF interactions were present in the TRMs we detected. For instance the pairwise TRM of ATF2-JUND(Log_2_NID score = 2.57) where both the TFs are part of the well-known AP-1 factor complex, which is involved in expression regulation of multiple TGs[30]–[32]. TF GATAD2B is known to form a repressive complex involving nucleosome remodeling and deacetylase activity with the CHD family of TFs[33]. We discovered that GATD2B and CHD1 were present in two different TRMs: ARID3A-CHD1-GATAD2B(Log_2_NID score = 2.64) and ARID3A-CHD1-GATAD2B-RELA(Log_2_NID score = 2.63). The presence of ARID3A and RELA in these TRMs has not been validated by the existing literature, although both of them have been associated with immune cell proliferation[34], [35]. We also discovered the three way TRM EZH2-KDM5A-SUZ12(Log_2_NID score =2.40, where the methyltransferase EZH2 and scaffolding protein SUZ12 are known to form the polycomb-repressive complex PRC2, which interacts and competes with H3K4me3 demethylase KDM5A during the process of angiogenesis and hematopoiesis[36]. We also discovered the pairwise TRM KDM5A-SUZ12(Log_2_NID score = 2.47) indicating that KDM5A and SUZ12 may be the primary interactors within the three-way TRM.

We also detected several TRMs containing previously uncharacterized TF interactions. For example, the TRM with the highest influence over TG expression was RELB-STAT1 with the largest Log_2_NID score of 3.18. Both of these TFs play an important role in immune response and lymphocyte development [37], [38]. Thus, their closely related functions could point to the possibility of their interaction *in vivo*. Another intriguing, albeit unvalidated interaction, that we discovered was EP300-TAF1(Log_2_NID score = 2.39). Both of these TFs are well known lysine acetyltransferases and are responsible for activating and regulating transcription of several TGs and were also found to have the highest frequency of oncogenic mutations among all the other lysine acetyltransferases[39]. The Log_2_NID scores for all pairwise TRMs are shown in the form of a heat-map in the ***Figure 2C*** (all scores are available in ***Supplementary table S3B***). Thus, we detected TRMs containing many previously uncharacterized as well as some well-known TF interactions using the NID algorithm.

### Chromatin looping plays an essential role in forming transcription factor regulatory modules and in mediating their regulation of target genes

Apart from some well-known interactions, our discovered TRMs contained a significant number of previously uncharacterized TF interactions. As described in the **Background**, TRM formation can be brought about by either co-localization of proximally binding TFs based on motif proximity or by distally binding TFs brought in close proximity by long distance chromatin looping. Thus, to characterize the TRMs, we used chromatin looping and TF motif co-occurrence information to identify TFs interacting with each other via chromatin looping (long-distance interactions) or by binding in close proximity(see **Detecting co-binding TF ChIP-Seq peaks** and **Detecting TF ChIP-Seq peaks interacting via chromatin looping** sections of the **Methods**).

Using GM12878 specific high throughput chromatin capture(Hi-C) data, we looked for long distance interactions between ChIP-seq peaks, present within TG’s *cis*-regulatory regions, corresponding to all pairwise TF modules we detected (***Figure 3A***). We compared the enrichment of Hi-C contacts for these peaks with that obtained from a background set of peak pairs, within the TG’s *cis*-regulatory regions, corresponding to random pairwise combinations of TFs not present in the detected set of pairwise TRMs using a chi-square test (see ***Supplementary table S4A***). We observed significant enrichment of Hi-C contacts at 5Kb resolution among 36,734 ChIP-seq peak pairs corresponding to 31 pairwise TF modules (χ^2^ p-value = 9e-04) within TG’s *cis* regulatory region as shown in ***Figure 3B*** and provided in ***Supplementary table S4B***. The only pairwise TRM that did not contain any Hi-C contact points between the peak pairs was SUZ12-ZNF284. The enrichment of Hi-C contacts at 1Kb resolution was not statistically significant, however with the χ^2^ p-value of 0.3423(see ***Supplementary Figure S3***).

**Figure 3:**
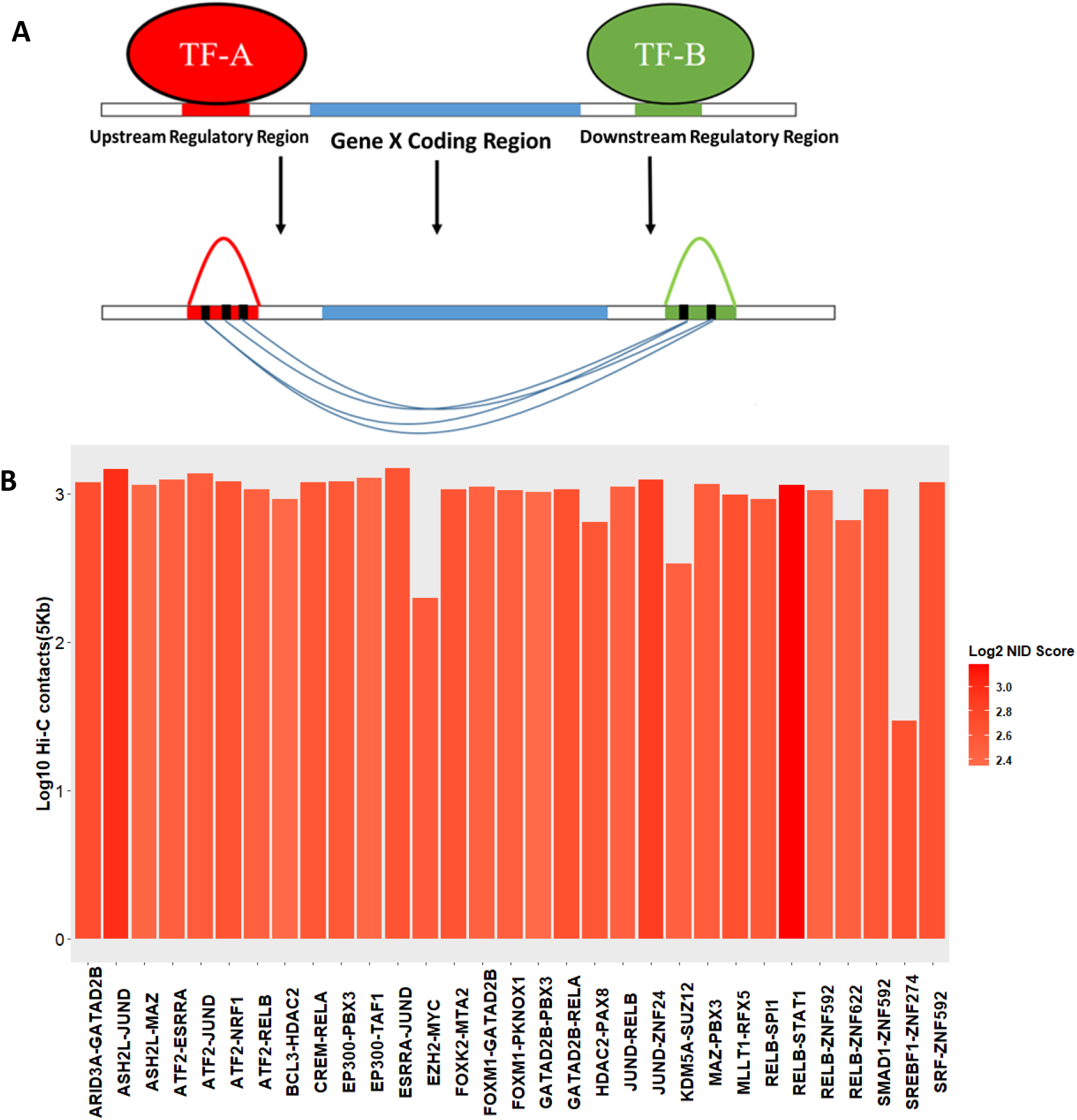
Pairwise TRMs interact via long distance chromatin looping. A) We overlapped the GM12878 Hi-C data at 5Kb resolution with the ChIP-seq peak pair regions corresponding to the 32 pairwise TRMs within the cis-regulatory regions of the TGs. B)Barplot showing the mean log10 Hi-C contacts(5Kb resolution) between peak regions of the pairwise TRMs shaded according to the respective Log2 NID scores across all the TG. We weren’t able to detect any HiC contacts between the peak pairs of the TRM SUZ12-ZNF284

In order to identify co-localizing TF interactions based on their sequence/motifs, we used the SpaMo tool from the MEME suite(*v.5.1.1*)[8] to examine pairwise TRMs,. We looked for significant spacing between TF motifs occurring within their overlapping peak pair regions. We found significant motif co-occurrence for 60 peak pairs corresponding to 6 pairwise modules (adjusted p-value < 0.05, see ***Supplementary table S4C***). Additionally, we did not find these co-binding TRMs in the set of modules previously described by other approaches[6], [7], [9], [40].

To further characterize the regulatory architecture of the TRMs, we defined four transcription regulatory programs shown in ***Figure 4A*** based on their interactions with TG promoters. We first identified 2,038 TGs where TFs peaks were interacting with each other either via Hi-C or via motif co-occurrence. We then determined the regulatory programs followed by the TRM peak pairs for each TG (see ***Supplementary table S5***). As shown in ***Figure 4B***, on an average 95% of the peak pairs corresponding to each pairwise TRM followed a configuration where at least one is interacting with the TG promoter and the two peaks interact with each other via long distance chromatin looping. Furthermore, TRMs HDAC2-PAX8, KDM5A-SUZ12 and SREBF1-ZNF274, for which the TFs are not known to directly bind to the TG promoters, regulated all their TGs using this program exclusively. We observed that for the remaining TRMs, about 4.5% of the peak pairs followed the second regulatory program which constituted one of them being present directly within the TG promoter while interacting with the other one via chromatin looping. About 17% of the peak pairs corresponding to the TRM EZH2-MYC, which contained TFs with known TG promoter binding activity, followed this regulatory program. Lastly, we found only 25 co-localizing peak pairs corresponding to 4 pairwise modules (RELB-SPI1, JUND-RELB, FOXM1-PKNOX1 and MAZ-PBX3) interacting with the promoters of 15 TGs via chromatin looping and 1 instance of co-localizing peak pair for the TRM RELB-SPI1 directly binding the promoter of 1 TG. Hence, only RELB-SPI1, which contained TFs important for lymphocyte development, contained peak pairs following all four types of transcription regulatory programs.

**Figure 4:**
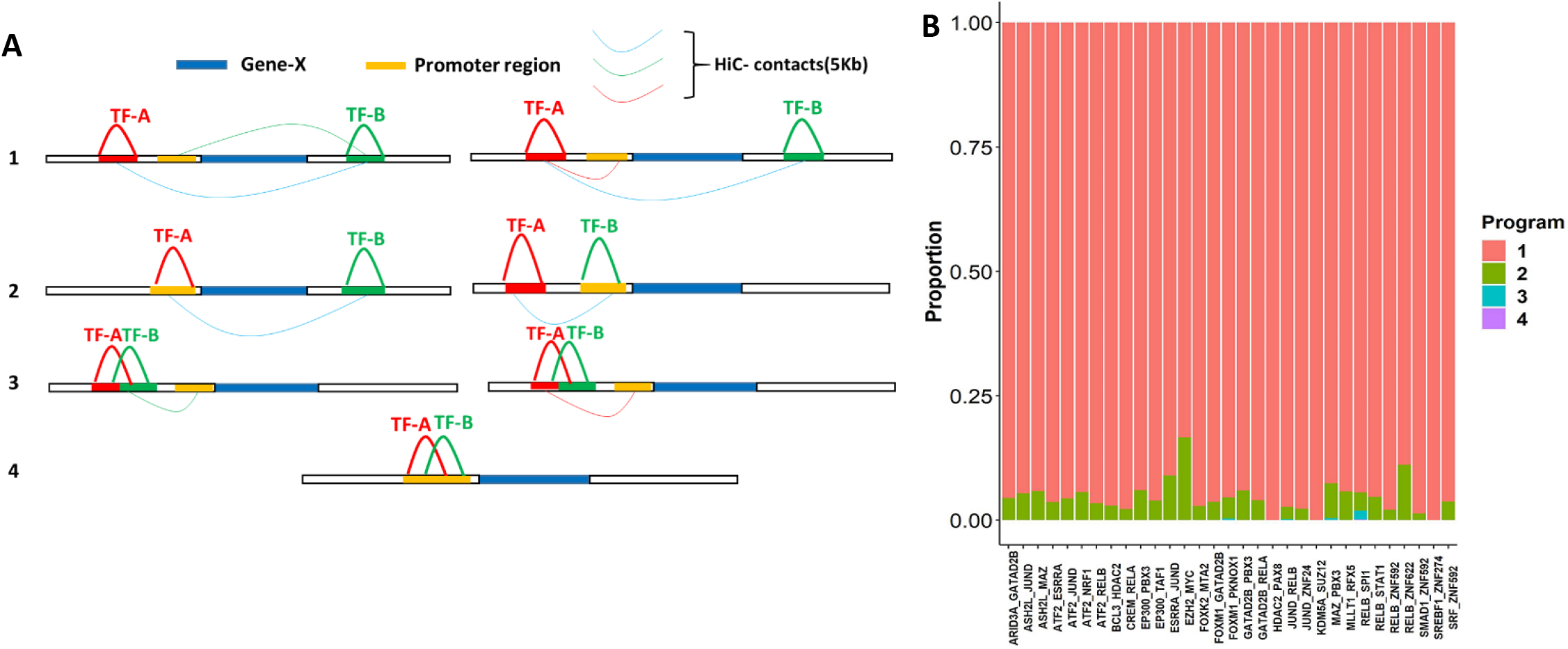
Pairwise TF TRMs follow different regulatory programs for different TGs. A) We utilized HiC and co-binding data to define 4 TF regulatory patterns/programs for the pairwise modules for different TGs. B) Barplot shows the proportion of the total peak pairs for each pairwise TRM following each of the 4 transcription regulatory programs shown in A

Thus, based on the above analyses, we conclude that the pairwise TRMs identified from the MLP-U learned models almost exclusively contained TF peak interactions occurring over long distance via chromatin looping. In addition, these TRMs mostly regulated their TGs also via long distance chromatin interactions with the TG promoters.

## Discussion

In this study, we designed a machine learning prediction framework for identifying TRM for the GM12878 immortalized LCL utilizing multiple big “omics” data sources. We used a modified form of the neural network MLP architecture called MLP-U in order to account for the influence of individual TFs as well as of TF interactions on TG expression within the same model. We found that accounting for both these effects resulted in more accurate TG expression prediction compared to accounting for just the linear effects of TFs using the ENET regularized regression models. The traditional MLP models produced better prediction than the MLP-U models because of the recapitulation of the main effects of TFs. In other words, both main effects and interaction effects were being modelled using complex non-linear functions in the traditional MLP architecture leading to perhaps an overestimation of the main effects resulting in the better TG expression prediction.

One of the biggest drawbacks of a neural network model is that it is usually considered a “black-box” as features learned during the training as well as testing of the models are difficult to interpret. We overcome this limitation and extracted biologically relevant information using the NID algorithm[29]. We calculated main effects of individual TFs as well as interaction effects of TF combinations. We observed that the direction of the TF main effects correlated well with their known functional roles. However, these effects were largely different compared those obtained from ENET models as the MLP-U captured context/interaction dependent TF main effects, while the ENET models estimate TF main effects only.

Furthermore, we also detected highly influential TF interactions forming TRMs via statistical interactions in models of TG expression. We derived literature-based annotations for some of these TRMs, while many were novel TF interactions not identified by other approaches. This could be due to two reasons. First, the non-additive non-linear nature of the TF interactions, reflecting the “TF collective model” we detected is fundamentally different from that of the linear, co-localizing TFs, reflecting the “Enhanceosome” and “Billboard” models identified by the previous approaches. Second, our strategy for identifying TF interactions was to model their influence on TG expression, which was largely ignored by the previous approaches. Thus, a co-localizing set of TFs not significantly impacting TG expression would be missed using our approach, though these TFs presumably have little influence on the expression of nearby genes. Additionally, we found that a significant proportion of the TF peaks for the pairwise TRMs interacted with each other and with the promoters of the TGs they regulated via chromatin looping. Therefore, long distance chromatin interactions likely play a large role in formation of TRMs as well as in their regulation of the TGs. This further validates the idea that TF interactions are not limited to proximally binding co-localizing sets of TFs. We used Hi-C chromatin looping data in two mutually independent contexts; we first included Hi-C contacts made by distal TFs with the TG promoters while building our GRNs, and further validated these chromatin interactions by examining Hi-C contact enrichments between the TF peak regions themselves. While the former Hi-C data aggregation was done to quantify the influence of distally binding TFs on TG regulation via promoter interaction, the latter instance reflected characterization of pairwise TFs interacting over long distances.

We focused our analyses on the GM12878 LCL in this study due to the density of TF binding data available, however our approach is flexible enough to analyze TRM based TG regulation in other commonly studied human cell-lines when these data are available A key limitation of our approach is the need for high-density omics assay data that often require large input DNA quantities that likely limit their application to cell-lines only. In different cellular contexts and environmental conditions, additional higher order TRMs may exist, and the precise models underlying these interactions will be difficult to elucidate. However, we did identify pairwise TF interactions that form a basis for higher order interactions that could act as a starting point for further experimental validation or examination under different environmental conditions.

## Conclusions

In this study, we have detected TRMs significantly impacting TG expression using neural network based prediction models containing multi-omics GRN derived TF regulatory features. We have demonstrated multiple ways in which long distance chromatin looping plays a role in TRM based TG regulation. Our approach for detection, characterization and validation of TRMs provides a roadmap for a multi-omics analysis to study the complex phenomenon of transcription regulation genome-wide, and may provide insights into the impact of transcriptional dysregulation in the genetic basis of human phenotypes.

## Methods

All the published algorithms and datasets used in this study have been described in ***Supplementary information***.

### Building multi-omics GRN

We utilized the Passing Attributes between Networks for Data Assimilation (PANDA) algorithm to build the GRN. This algorithm uses a TF binding site(TFBS) based motif network, a PPI network and a co-expression network for building the GRN(***Figure-1***). We generated these three networks using the following approach:

Motif network: We isolated all the ChIP-Seq peaks within a 50Kb window upstream of the TSS of the longest transcript and downstream of the body of each protein coding TG. We then used the most distant CTCF peaks to demarcate the *cis*-regulatory boundaries for these TFBS, as it is a well-known insulator protecting the enhancers of TG gene from acting upon the promoters of another as shown in ***Figure-1***. Furthermore, we added the TFBS found in the intronic regions of each TG to this set in order to capture the effect of introns on transcriptional regulation. We have shown previously that inclusion of intronic TFBS in the GRN framework ultimately improves the model prediction accuracy[41], as introns are hypothesized to have regulatory influence over TG expression[3], [42]. We then weighted each TFBS based on the number of Hi-C contacts(1Kb) it makes with the TG’s *promoter(**Figure-1***) using the weighting scheme described in **Generating Hi-C Weightings** of the ***Supplementary Methods***. Using such a weighting scheme helps to capture regulatory information provided by long distance interactions of distal TFBS with TG promoters created via chromatin looping while preserving the influence of proximal promoter based TFBS[41]. We created a weighted motif network using the unique TF-TG interactions and the average Hi-C weight for them.
PPI network: We downloaded PPI data from the BioGRID database(v.3.5.188) to generate the PPI network.
Co-expression network: We extracted expression residuals for the 462 LCL samples within the GEUVADIS datasets using a genome-wide genetic relationship matrix(GRM) based mixed-linear regression model and used them to generate the co-expression network (see **Building co-expression network for the PANDA GRN** of ***Supplementary Methods***). This was done to adjust out the genetic effect of the variants in the dataset.

We used the above networks to generate GRN utilizing the R(v.3.4.2) implementation of the PANDA algorithm. After 25 iterations, we obtained convergence by setting the threshold for Hamming’s distance at 0.001 and by using the value of 0.1 for the update parameter.

### MLP network architecture and building the prediction models

We utilized two different MLP architectures in our paper: 1)MLP-U(MLP-Univariate) and 2)Traditional MLP as shown in ***Supplementary Figure S1***. The MLP-U architecture contained individual univariate MLPs receiving inputs corresponding to each TF in addition to the traditional MLP. All the univariate MLPs had 3 layers containing 10 nodes each and the traditional MLP also contained 3 layers with 800, 500 and 1000 nodes for each model. The non-linear activation function for all the layers was Rectified Linear Unit(ReLU).

We built the ENET and the MLP prediction models using log10 FPKM expression values of 11,780 protein coding TGs, where we used 80% of the data(9,424 TGs) for training the models and the remaining 20%(2356 TGs) to test the models and assess their prediction accuracy. We used two different internal cross-validation strategies to train the two types of models: 1) For the MLP-U and MLP models, we further divided the training data into 85% training and 15% validation sets. We then trained these models using the backpropagation algorithm. Additionally, we summed the output from all the individual univariate MLPs and the traditional MLP at the last node for training the MLP-U models. We note here that the traditional MLP architecture was only used as a comparison in the paper and most of the analyses were done using the trained MLP-U models. 2) For the ElasticNet(ENET) prediction model, we used an alpha of 0.5 and trained the models based on 20 fold inner cross-validation. We trained and tested the models for 20 iterations(***Figure-1***), and computed Pearson’s Correlation Coefficient(PCC) each time to assess model performance.

Thus, we had an input matrix **X** of size N × T, containing N TGs and T TFs. The values in this matrix were scaled edge-weights corresponding to the vertex TF_t_ ⟶TG_n_, where n ϵ{N} and t ϵ {T} derived from the learned PANDA GRN network. The output was a column vector **y** of size *N* containing scaled and centered log FPKM (Fragments per kilobase per million) expression values of the *N* genes. For the MLP-U models, it was derived based on a generalized additive model:

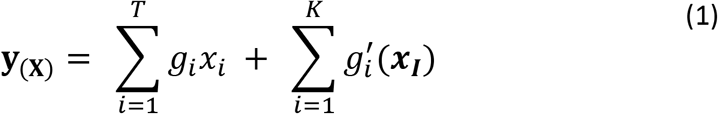

### Obtaining main and interaction effects from the MLP-U models

For each trained MLP-U model, we performed an additional 5-fold prediction task in order to capture the prediction performance over all the TGs within each iteration. Thus, we essentially conducted 100 prediction rounds for which we stored the model weights learned during the training process.

In order to calculate the main effect corresponding to each TF, we utilized the learned MLP-U models. Specifically, we extracted layer weights from each one of the univariate MLP corresponding to each TF feature and aggregated them across all the prediction iterations. These iterations corresponded to a set *S* of 20 random numbers *s* each representing an instance/state for bootstrapping test set genes for each prediction task.

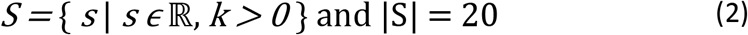

For each random state *s*, we picked 5 non-overlapping sets of test genes

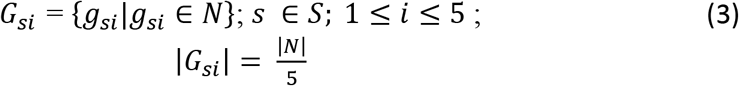

For each *G_si_*, we then used the remaining genes as the training set *G_si_train_* such that

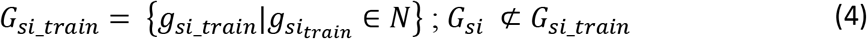

We then predicted the expression values of *G_si_* genes according to the following equation:

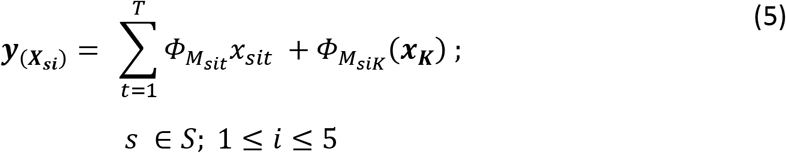

Here **Y**(*X_si_*) is the vector containing predicted expression for gene set *G_si_* using the input matrix ***X_si_*** by the model trained using the input from genes in set *G_si_train_*. The first part of equation (5) captures the main effect of each one of the TF *t* with *M_sit_* representing the corresponding univariate MLP while the second part captures the interaction effect of *K* interactions, via the traditional MLP *M_sik_*, on the gene expression trait. Thus, for each iteration s, the expression vector of gene set *G_si_* ***Y***(*X_si_*) is derived from a generalized additive model *M_si_* containing main effects and interaction effects derived from a collection of complex non-linear functions Φ*M_sit_* and *ΦM_siν_* respectively. The parameters for this model were learned during the training process using the training set *G_si_train_*. Furthermore, we had 5 models for each random iteration each containing a different set of test genes.

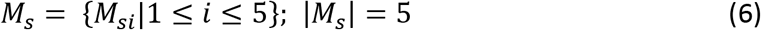

The architecture for each model, w.r.t the number of hidden units in each layer and the number of hidden layers was similar. Each model *M_sit_* and *M_siν_* contained *L* hidden layers, and there were *p_l_* units/neurons in the *l*-th layer. The input layer for the univariate MLP *M_sit_* was the vector ***x**_sit_* containing edge-weights for TGs corresponding to TF *t* 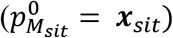. On the other hand, the input layer for the traditional MLP *M_siκ_* was the matrix ***X_si_*** containing the edgeweights corresponding to all the 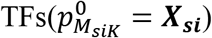. In each model, there were ***L*** weight matrices containing the weights learned during the training process such that 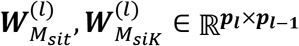, *l* = 1,2,. *L* and *L* + 1 bias vectors 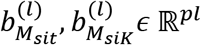, *l* = 0,1,2.*L*. Furthermore, there is a non-linear activation function *ϕ*(·) associated with each unit and weights 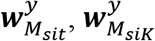 and biases 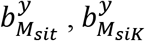 associated with the output layer for each model. The hidden units 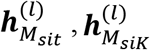 and the outputs 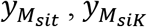 for the models can be mathematically described as:

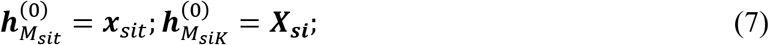

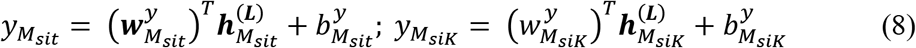

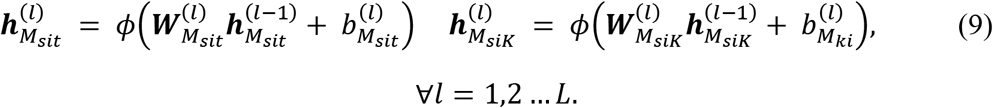

We note here that the ***L*** = 3 for all the models in our case.

We utilized the learned models *M_sit_* and *M_siκ_* to calculate the main effect for each TF *t* and the interaction effect of *K* interactions respectively. We used an extension of the neural interaction detection(NID) developed by Tsang *et al*. in order to compute these effects[29].

For each random state s, we first aggregated the layer weights across all the models

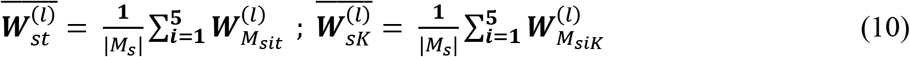

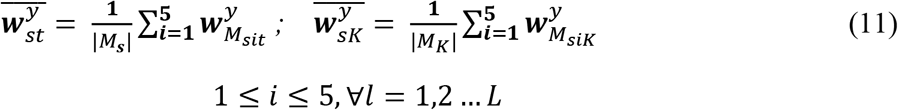

Here, 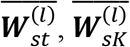 and 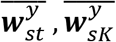 represent the weights of each hidden layer and the output layers respectively averaged across all the models in *M_s_*.

The main effect for each TF *t* and the interaction effect of the *TF_m_-TF_n_* interaction at unit *j* of the first layer across all the models for a random state s was calculated using the following equations:

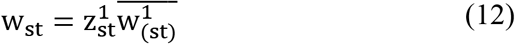

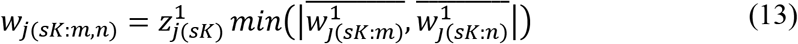

Here, W_(st)_ is the main effect of the transcription factor t obtained from the first layer of univariate model corresponding to random state s, 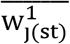 is the mean weight of all the connections made by the input node in the first layer. Similarly, Wj_(sK:m,n)_ is the interaction effect for the interaction between *TF_m_* and *TF_n_* at the hidden unit *j* of the first layer aggregated across all the models in *M_s_* and 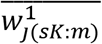 and 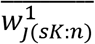 are the aggregated weights corresponding to the connections(indices) of *TF_m_* and *TF_n_* respectively at node *j*. 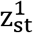 and 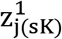 represent the influence of the input node and the hidden unit *j* respectively, which are calculated using the following formulae:

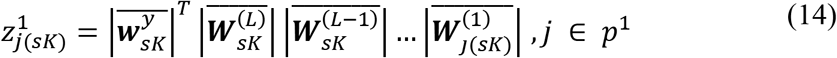

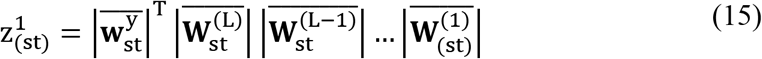

The aggregated weight of interaction between *TF_m_* and *TF_n_* across all the nodes in the first layer was calculated using the following equation:

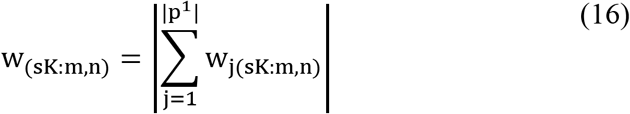

This step was not necessary for the main effects calculation since we only had one input node in each univariate MLP corresponding to each TF.

Since we averaged the calculations over all the models that contained different sets of test genes for each random state, we assumed that w_(sν:m,n)_ and w_st_ represented average interaction effect between *TF_m_* and *TF_n_* and average main effect of TF t respectively over all the genes. We then averaged this effect over all the random states to produce the final NID interaction effects and main effects:

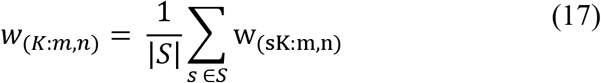

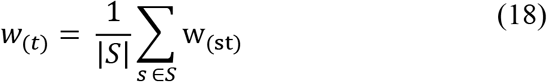

### Calculating TF average ENET main effects

We calculated the average effect estimate for TF *T* 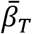 using the following equation:

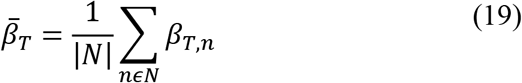

Here, *N* is the set of random instances that we used to build our ENET prediction models and *β_T,n_* is the effect estimate of *T* for instance *n*. We calculated these effect estimates for each one of the 149 TFs and tabulated them in ***Supplementary table S3B***.

### Detecting co-binding TF ChIP-Seq peaks

In order to identify statistically significantly co-binding pairs of TF ChIP-Seq peaks, we utilized the SpaMo algorithm (meme suite version 5.1.1)[8], which looks for significantly enriched spacings between a primary motif and a secondary motif by within a set of sequences. We isolated all the overlapping peak pairs corresponding to the 32 pairwise TF modules present within the TG’s *cis*-regulatory regions. We centered and modified these regions so that they are no longer than 500bp, which is the required size for sequences for SpaMo. We utilized the position weight matrices(PWMs) downloaded from HOCOMOCO(*v.11*)and JASPAR(*v.2020*) in order to scan the sequences for motifs corresponding to TFs in each pairwise TRM. We ran the SpaMo command line version and extracted peak pairs representing co-localizing TFs at a p-value threshold of 0.05.

### Detecting TF ChIP-Seq peaks interacting via chromatin looping

We used the Hi-C data downloaded for GM12878(GEO accession: GSM1551688) in order to look for TF peaks interacting via chromatin looping. We used data corresponding to 1Kb and 5Kb resolution, and overlapped the peak pairs of pairwise TRMs with the Hi-C contact points. We also generated a random set of peak pairs corresponding to pairwise TFs not forming a pairwise TRM representing the background set for performing χ^2^ test of enrichment (see ***Supplementary Table S4B***). We tested for enrichment of Hi-C contacts within the peak pairs corresponding to the TRMs detected using this test at a p-value threshold of 0.05.

## List of abbreviations

TF: Transcription Factors
TG: Target Gene
ChIP-Seq: Chromatin Immunoprecipitation Sequencing
PANDA: Passing Attributes between Networks for Data Assimilation
GRN: Gene Regulatory Network
TFBS: Transcription Factor Binding Site/s
ENET: ElasticNet
FPKM: Fragments per kilobase of transcripts per million
PCC: Pearsons Correlation Coefficient
PPI: Protein-protein interactions
ENCODE: Encyclopedia of DNA elements
IDR: Irreproducible Discovery Rate
TRM: TF regulatory module
MLP: Multilayer Perceptron
MLP-U: Univariate MLPs with traditional MLP
SpaMo: Spaced motif analysis
LCL: Lymphoblastoid cell line
NID: Neural Interaction Detection

## Declarations

### • Ethics approval and consent to participate

The study makes use of publicly accessible data.

### • Consent for publication

Not applicable

### • Availability of data and materials

All the datasets used in this paper are publicly available. The code used in the project is posted on github (https://github.com/bushlab-genomics/TRM-Detection).

### • Consent for publication

Not applicable

### • Authors’ contributions

NP and WSB conceptualized and designed the research project, and drafted the manuscript. NP collected the data and performed all subsequent analyses.

## References

[1] S. A. Lambert et al., “The Human Transcription Factors,” Cell, vol. 172, no. 4, pp. 650–665, 2018, doi: https://doi.org/10.1016/j.cell.2018.01.029.

[2] L. Prazak, M. Fujioka, and J. P. Gergen, “Non-additive interactions involving two distinct elements mediate sloppy-paired regulation by pair-rule transcription factors,” Dev. Biol., vol. 344, no. 2, pp. 1048–1059, Aug. 2010, doi: 10.1016/j.ydbio.2010.04.026.

[3] J. I. Fuxman Bass et al., “Transcription factor binding to Caenorhabditis elegans first introns reveals lack of redundancy with gene promoters,” Nucleic Acids Res., vol. 42, no. 1, pp. 153–162, Jan. 2014, doi: 10.1093/nar/gkt858.

[4] F. Spitz and E. E. M. Furlong, “Transcription factors: from enhancer binding to developmental control,” Nat. Rev. Genet., vol. 13, no. 9, pp. 613–626, 2012, doi: 10.1038/nrg3207.

[5] C. M. Vockley, I. C. McDowell, A. M. D’Ippolito, and T. E. Reddy, “A long-range flexible billboard model of gene activation,” Transcription, vol. 8, no. 4, pp. 261–267, Aug. 2017, doi: 10.1080/21541264.2017.1317694.

[6] M. B. Gerstein et al., “Architecture of the human regulatory network derived from ENCODE data,” Nature, vol. 489, no. 7414, pp. 91–100, 2012, doi: 10.1038/nature11245.

[7] Y. Guo and D. K. Gifford, “Modular combinatorial binding among human trans-acting factors reveals direct and indirect factor binding,” BMC Genomics, vol. 18, no. 1, p. 45, 2017, doi: 10.1186/s12864-016-3434-3.

[8] T. Whitington, M. C. Frith, J. Johnson, and T. L. Bailey, “Inferring transcription factor complexes from ChIP-seq data,” Nucleic Acids Res., vol. 39, no. 15, pp. e98–e98, May 2011, doi: 10.1093/nar/gkr341.

[9] G. Yang, A. Ma, Z. S. Qin, and L. Chen, “Application of topic models to a compendium of ChIP-Seq datasets uncovers recurrent transcriptional regulatory modules,” Bioinformatics, vol. 36, no. 8, pp. 2352–2358, Jan. 2020, doi: 10.1093/bioinformatics/btz975.

[10] K. Izumi, “Disorders of Transcriptional Regulation: An Emerging Category of Multiple Malformation Syndromes,” Mol. Syndromol., vol. 7, no. 5, pp. 262–273, 2016, doi: 10.1159/000448747.

[11] T. I. Lee and R. A. Young, “Transcriptional Regulation and Its Misregulation in Disease,” Cell, vol. 152, no. 6, pp. 1237–1251, Mar. 2013, doi: 10.1016/j.cell.2013.02.014.

[12] K. R. Pennypacker, “AP-1 transcription factor complexes in CNS disorders and development.,” J. Fla. Med. Assoc., vol. 82, no. 8, pp. 551–554, Aug. 1995.

[13] S. Trop-Steinberg and Y. Azar, “AP-1 Expression and its Clinical Relevance in Immune Disorders and Cancer.,” Am. J. Med. Sci., vol. 353, no. 5, pp. 474–483, May 2017, doi: 10.1016/j.amjms.2017.01.019.

[14] A. Mortazavi et al., “Integrating and mining the chromatin landscape of cell-type specificity using self-organizing maps,” Genome Res., vol. 23, no. 12, pp. 2136–2148, Dec. 2013, doi: 10.1101/gr.158261.113.

[15] Z. Yang and G. Michailidis, “A non-negative matrix factorization method for detecting modules in heterogeneous omics multi-modal data,” Bioinformatics, vol. 32, no. 1, pp. 1–8, Sep. 2015, doi: 10.1093/bioinformatics/btv544.

[16] E. G. Giannopoulou and O. Elemento, “Inferring chromatin-bound protein complexes from genome-wide binding assays,” Genome Res., vol. 23, no. 8, pp. 1295–1306, Aug. 2013, doi: 10.1101/gr.149419.112.

[17] B. P. Berman et al., “Exploiting transcription factor binding site clustering to identify cis-regulatory modules involved in pattern formation in the Drosophila genome,” Proc. Natl. Acad. Sci. U. S. A., vol. 99, no. 2, pp. 757–762, Jan. 2002, doi: 10.1073/pnas.231608898.

[18] K. Glass, C. Huttenhower, J. Quackenbush, and G.-C. Yuan, “Passing Messages between Biological Networks to Refine Predicted Interactions,” PLoS One, vol. 8, no. 5, p. e64832, May 2013.

[19] N. Patel and W. S. Bush, “Modeling transcriptional regulation using gene regulatory networks based on multi-omics data sources,” BMC Bioinformatics, vol. 22, no. 1, p. 200, 2021, doi: 10.1186/s12859-021-04126-3.

[20] A. C. Vaqueiro et al., “Expanding the spectrum of TBL1XR1 deletion: Report of a patient with brain and cardiac malformations,” Eur. J. Med. Genet., vol. 61, no. 1, pp. 29–33, 2018, doi: https://doi.org/10.1016/j.ejmg.2017.10.008.

[21] S. Gordon, G. Akopyan, H. Garban, and B. Bonavida, “Transcription factor YY1: structure, function, and therapeutic implications in cancer biology,” Oncogene, vol. 25, no. 8, pp. 1125–1142, 2006, doi: 10.1038/sj.onc.1209080.

[22] G. Rodier et al., “The Transcription Factor E4F1 Coordinates CHK1-Dependent Checkpoint and Mitochondrial Functions,” Cell Rep., vol. 11, no. 2, pp. 220–233, Apr. 2015, doi: 10.1016/j.celrep.2015.03.024.

[23] C. S. Hill, “Transcriptional Control by the SMADs,” Cold Spring Harb. Perspect. Biol., vol. 8, no. 10, p. a022079, Oct. 2016, doi: 10.1101/cshperspect.a022079.

[24] M. de Dieuleveult and B. Miotto, “DNA Methylation and Chromatin: Role(s) of Methyl-CpG-Binding Protein ZBTB38,” Epigenetics insights, vol. 11, pp. 2516865718811117–2516865718811117, Nov. 2018, doi: 10.1177/2516865718811117.

[25] S. Ropero, E. Ballestar, M. Alaminos, D. Arango, S. Schwartz, and M. Esteller, “Transforming pathways unleashed by a HDAC2 mutation in human cancer,” Oncogene, vol. 27, no. 28, pp. 4008–4012, 2008, doi: 10.1038/onc.2008.31.

[26] T. Ismail, H.-K. Lee, C. Kim, T. Kwon, T. J. Park, and H.-S. Lee, “KDM1A microenvironment, its oncogenic potential, and therapeutic significance,” Epigenetics Chromatin, vol. 11, no. 1, p. 33, 2018, doi: 10.1186/s13072-018-0203-3.

[27] L. Icardi et al., “The Sin3a repressor complex is a master regulator of STAT transcriptional activity,” Proc. Natl. Acad. Sci., vol. 109, no. 30, pp. 12058 LP–12063, Jul. 2012, doi: 10.1073/pnas.1206458109.

[28] W. Akhtar and G. J. C. Veenstra, “TBP-related factors: a paradigm of diversity in transcription initiation,” Cell Biosci., vol. 1, no. 1, p. 23, 2011, doi: 10.1186/2045-3701-1-23.

[29] M. Tsang, D. Cheng, and Y. Liu, “Detecting Statistical Interactions from Neural Network Weights.” 2017.

[30] A. Giannoudis et al., “Activating transcription factor-2 (ATF2) is a key determinant of resistance to endocrine treatment in an in vitro model of breast cancer,” Breast Cancer Res., vol. 22, no. 1, p. 126, 2020, doi: 10.1186/s13058-020-01359-7.

[31] E. Shaulian and M. Karin, “AP-1 in cell proliferation and survival,” Oncogene, vol. 20, no. 19, pp. 2390–2400, 2001, doi: 10.1038/sj.onc.1204383.

[32] J. M. Hernandez, D. H. Floyd, K. N. Weilbaecher, P. L. Green, and K. Boris-Lawrie, “Multiple facets of junD gene expression are atypical among AP-1 family members,” Oncogene, vol. 27, no. 35, pp. 4757–4767, 2008, doi: 10.1038/onc.2008.120.

[33] C. Shieh et al., “GATAD2B-associatedneurodevelopmental disorder (GAND): clinical and molecular insights into a NuRD-relateddisorder,” Genet. Med., vol. 22, no. 5, pp. 878–888, 2020, doi: 10.1038/s41436-019-0747-z.

[34] H. L. Pahl, “Activators and target genes of Rel/NF-κB transcription factors,” Oncogene, vol. 18, no. 49, pp. 6853–6866, 1999, doi: 10.1038/sj.onc.1203239.

[35] J. M. Ward et al., “Human effector B lymphocytes express ARID3a and secrete interferon alpha,” J. Autoimmun., vol. 75, pp. 130–140, Dec. 2016, doi: 10.1016/j.jaut.2016.08.003.

[36] J. Chen et al., “Two faces of bivalent domain regulate VEGFA responsiveness and angiogenesis,” Cell Death Dis., vol. 11, no. 1, p. 75, 2020, doi: 10.1038/s41419-020-2228-3.

[37] M. Yang et al., “Biological characteristics of transcription factor RelB in different immune cell types: implications for the treatment of multiple sclerosis,” Mol. Brain, vol. 12, no. 1, p. 115, 2019, doi: 10.1186/s13041-019-0532-6.

[38] C. K. Lee, E. Smith, R. Gimeno, R. Gertner, and D. E. Levy, “STAT1 affects lymphocyte survival and proliferation partially independent of its role downstream of IFN-gamma.,” J. Immunol., vol. 164, no. 3, pp. 1286–1292, Feb. 2000, doi: 10.4049/jimmunol.164.3.1286.

[39] Y. Jiang et al., “Metagenomic characterization of lysine acetyltransferases in human cancer and their association with clinicopathologic features,” Cancer Sci., vol. 111, no. 5, pp. 1829–1839, May 2020, doi: 10.1111/cas.14385.

[40] Y. M. Oh, J. K. Kim, S. Choi, and J.-Y. Yoo, “Identification of co-occurring transcription factor binding sites from DNA sequence using clustered position weight matrices,” Nucleic Acids Res., vol. 40, no. 5, pp. e38–e38, Dec. 2011, doi: 10.1093/nar/gkr1252.

[41] N. Patel and W. Bush, “No Title,” BMC Bioinformatics, 2021, doi: 10.21203/rs.3.rs-112300/v1.

[42] A. B. Rose, “Introns as Gene Regulators: A Brick on the Accelerator,” Frontiers in Genetics, vol. 9. p. 672, 2019.

