## Supplementary File for "Detecting global influence of transcription factor interactions on gene expression in lymphoblastoid cells using neural network models"

**Datasets used in the study**

**ChIP-Seq Files**

In order to obtain information about the TFBS across genomes for GM12878, we used the Encyclopedia of DNA elements(ENCODE) database[1]. We downloaded processed ChIP-seq narrow peak bedfiles corresponding to 149 TFs corresponding to GM12878 LCL that were aligned with hg19/GRCh37 reference assembly of the human genome and that had passed the optimal IDR threshold as defined by the ENCODE consortium. The accession number for these files are provided in ***Supplementary table S1*.**

**Gene Annotations**

We used the GRCh37/hg19 reference genome build from the biomaRt library(v.2.44.1)[2] in R to derive gene annotations such as transcription start sites(TSS), length of the gene body, transcript ids, exon ids, transcript lengths, gene ids etc. for all the protein coding genes.

**PWM Files**

We downloaded the position weight matrix(PWM) files corresponding to 469 human TFs from the JASPAR database(v.2020)[3] in MEME format. We later used these files as inputs for running the FIMO algorithm [4]in order to find statistically TFBS across the genome.

**TFBS sequences**

We used the GRCh37/hg19 reference build to obtain sequences corresponding the transcription factor binding sites(TFBS) in the regulatory region of each gene. We later used these sequences as inputs along with the PWMs for running the SpaMo algorithm[5] in order to find statistically significant spacings between overlapping TF peaks.

**Protein-Protein Interaction Data**

We used the BioGrid database(v.3.5.188)[6] to download PPI data in order to build the PANDA GRNs. We only used the high confidence experimentally validated using experimental techniques such as co-fractionation, co-immunoprecipitation, yeast two-hybrid and affinity capture in BioGrid. We further filtered out the PPIs that did not contain TFs, which ultimately provided us with 1937 PPIs among the GM12878 TFs.

**Co-expression data**

We used the GM12878 LCL expression data from the GEUVADIS project[7], which contains lymphoblastoid RNA-seq and genotype data derived from individuals belonging to European and African ancestry groups who participated in the 1000 genomes project, to build the co-expression network.

**Processing expression data for the prediction models**

We downloaded RNA-seq data for GM12878(ENCSR889TRN) from the ENCODE database, which contained processed TG quantification data for two technical replicates. We used the Log10 normalized mean FPKM values as outcome for the prediction models.

**Algorithms used in this study**

**PANDA**

Passing Attributes between Networks for Data Assimilation(PANDA) is a GRN building algorithm developed by Glass et al. to capture information from information corresponding to TF based regulatory mechanisms such as cooperativity among different combinations of TFs and co-regulation of multiple TGs by the same TF in order to weight the regulatory interactions between TF and TGs[8]. It iteratively updates the edge-weights of the regulatory network containing edges between TF and TG by using two Tanimoto similarity based metrics: Responsibility and Availability.

The responsibility of an edge is calculated using the information from the protein-protein interaction(PPI) network, while the availability is calculated from the gene co-expression network. Mathematically, the responsibility of an edge from TF $i$ to its TG $j$ for iteration t, ($R_{\mathrm{ij}}^{(t)}$) is calculated using the following equation:

$$R_{ij}^{(t)}= \frac{\sum_{m} P_{im}^{(t)}W_{mj}^{(t)}}{\sqrt{\sum_{m} \left( P_{im}^{(t)} \right)^{2}+ \sum_{m} \left( W_{mj}^{(t)} \right)^{2}- \left| \sum_{m} P_{im}^{(t)}W_{mj}^{(t)} \right|}}$$

Here, $P_{\mathrm{im}}^{(t)}$is the weight of the edge formed by TF $i$ with another TF $m$ in the PPI network at iteration $t$, while $W_{\mathrm{mj}}^{(t)}$is the edge weight for the connection between $m$and TG $j$in the regulatory network at iteration $t$. The agreement between the PPI and the regulatory network is thus calculated for each edge at each iteration using the cooperativity information among a set of TFs regulating the same set of TGs.

Similarly, the availability of an edge from TF $i$ to its TG $j$ for iteration t, ($A_{\mathrm{ij}}^{(t)}$) is calculated using the co-expression network based on the following equation:

$$A_{ij}^{(t)}= \frac{\sum_{k} W_{ik}^{(t)}C_{kj}^{(t)}}{\sqrt{\sum_{k} \left( W_{ik}^{(t)} \right)^{2}+ \sum_{k} \left( C_{kj}^{(t)} \right)^{2}- \left| \sum_{k} W_{ik}^{(t)}C_{kj}^{(t)} \right|}}$$

Here, $W_{\mathrm{ik}}^{(t)}$ represents the edge weight of the connection between TF $i$ and TG $k$ in the regulatory network and $C_{\mathrm{kj}}^{(t)}$ is the weight of the edge between TGs $k$ and $j$ in the co-expression network. Thus, this equation measures the availability of a TF based on the number of genes that it coregulates.

The weight of the edge between TF $i$ and TG $j$ at each iteration $t$ is then updated by averaging $A_{\mathrm{ij}}^{(t)}$and $R_{\mathrm{ij}}^{(t)}$ as well as a small update parameter $\alpha$:

$$\tilde{W}_{ij}^{(t)}= {0.5R}_{ij}^{(t)}+0.5A_{ij}^{(t)}$$

$$W_{ij}^{(t+1)}=\left( 1-\alpha\right)W_{ij}^{\left( t \right)}+ \alpha\tilde{W}_{ij}^{(t)}$$

Just as information is passed into the regulatory network from PPI and co-expression networks, information is passed out to the two networks using similar methods. The whole process is repeated and the edge-weights are updated until the network reaches convergence, which is determined using Hamming distance:

$$H= \left| \tilde{W}^{(t)}-W^{(t-1)} \right|= \frac{1}{N}\sum_{i,j} \left| \tilde{W}_{ij}^{(t)}- W_{ij}^{(t-1)} \right|$$

Here, $N$ is the number of possible edges in the regulatory network calculated by multiplying the number of TFs and TGs in the network. We used the R package pandaR to implement the PANDA algorithm in our project.

**Elastic-Net(ENET) regularized regression**

We used ENET regularized regression models, which linearly combine the lasso(L1) and the ridge(L2) penalty norms for feature selection and handling multi-colinearity among the features, for predicting gene expression.[9]

In the context of gene regulation, ENET uses a combination of two different penalizing methods to find the optimum number of regulators as predictors based on their influence on each other and on the expression of their target gene according the following equation:

$$\hat{\beta}=\arg\min_{\beta} \left\| y-X\beta\right\|+ \alpha\left\| \beta\right\|^{2}+(1- \alpha)\left\| \beta\right\|$$

(S1)

Here, $\hat{\beta}$ is the estimated effect coefficient for each input feature(TF), $\beta$ is the effect coefficient for each input feature, $X$ is the input feature matrix containing panda edge weights or tepic, affinity scores, $y$ is the response vector (expression values) and $\alpha$ is the hyperparameter used to control the ratio between the lasso and the ridge penalty norms.

**SpaMo**

Spaced motif analysis(SpaMo) was designed to identify significant spacing between motifs to two proximally binding co-localizing TFs[5]. It works on the hypothesis that if two TFs bind at a fixed distance in the given set of input DNA sequences then there is a high probability that they form a complex with each other. Working on this hypothesis, SpaMo scans a given set of DNA sequences for the presence of a primary motif corresponding to a TF and identifies hits based on position weight matrix scores. It then scans the sequences for the presence of a set of secondary motifs corresponding to a set of TFs and finds significant hits using position weight matrices again. It then calculated displacement between the primary motif and the set of secondary motifs and derives p-value using the null hypothesis that the displacements between a pair of primary and secondary will follow a uniform distribution if there is no significant interaction between the TFs. On the other hand, a binomial distribution of the displacement would reflect significant interaction between a pair of TFs.

**Supplementary Methods**

**Generating Hi-C Weightings**

We accessed Hi-C data GM12878(GSM1551688) with 1Kb and 5Kb resolution. We defined the promoter as the 5Kb region upstream of the TSS of the longest transcript for each gene. We normalized the Hi-C interactions using the Knight Ruiz(KR) normalization and created sparse contact matrices for both cell types. We calculated the number of contact points between each TF peak within a gene’s distal regulatory region and its promoter using *bedtools* *v.2.27.1*. We then calculated the HiC adjusted edge-weights between each TF and TG using the following formula:

$$C_{i,g}=1+ scaled(\frac{1}{N_{i,g}}\sum_{p\epsilon P_{i,g}} c_{p})$$

(S2)

Here, $C_{i,g}$ is the Hi-C adjusted edge weight between TF $i$ and TG $g$ , $N_{i,g}$ is the number of ChIP-seq peaks corresponding to $i$ in the regulatory region of $g$, $P_{i,g}$ is the set of peaks corresponding to $i$ in the regulatory region of $g$ and $c_{p}$ is the number of KR normalized contacts made by peak $p$ with the promoter of $g$. We used the MinMax scaling function of the *scikit-learn* library to scale the mean contacts within the (0,0.99) range. Thus, if the TF did not contain any peaks interacting with a gene’s promoter, the $C_{i,g}$ would be equal to 1 and the maximum value for $C_{i,g}$would be 1.99. We then extracted all the promoter-based TF-TG interactions that were down-weighted to 1.0, or were found to have no Hi-C interactions, and gave them maximum weight of 2.0 to create the cell-type specific “Hi-C UP” adjacency matrix**.**

**Building co-expression network for the PANDA GRN**

We used the Log normalized expression values(log FPKM) for the 15,785 protein coding genes from the lymphoblastoid cells of 462 individuals in the GEUVADIS dataset with variant effects regressed out using mixed-linear models with a genome-wide genetic relationship matrix(GRM). Our models could be described using the equation below:

  $y=X \beta+Zu+ \epsilon$

Here, y is the vector containing log FPKM expression values for the 462 individuals,  X is the matrix of size 462 by N , where N represents the number of common variants (minor allele frequency > 0.05) present in the dataset (6,326,925), containing the additive genotypes for each variant for each individual,  β is the vector of size N by 1 containing the effect estimates/coefficients of each variant obtained from the fitted regression models;  Z is the GRM of size 462 by 462   built using the number of alleles shared by each pair of individuals at the loci representing all the 6,326,925 variants across the genome; u is the random effects vector of size 462 capturing the random variance for each individual from the GRM and finally  ϵ is the residual vector of size 462 containing the effects not explained by the model. After fitting the models across all the genes, we extracted the ϵ term for each gene which contained the residual expression values. We used these values for building the co-expression matrix.

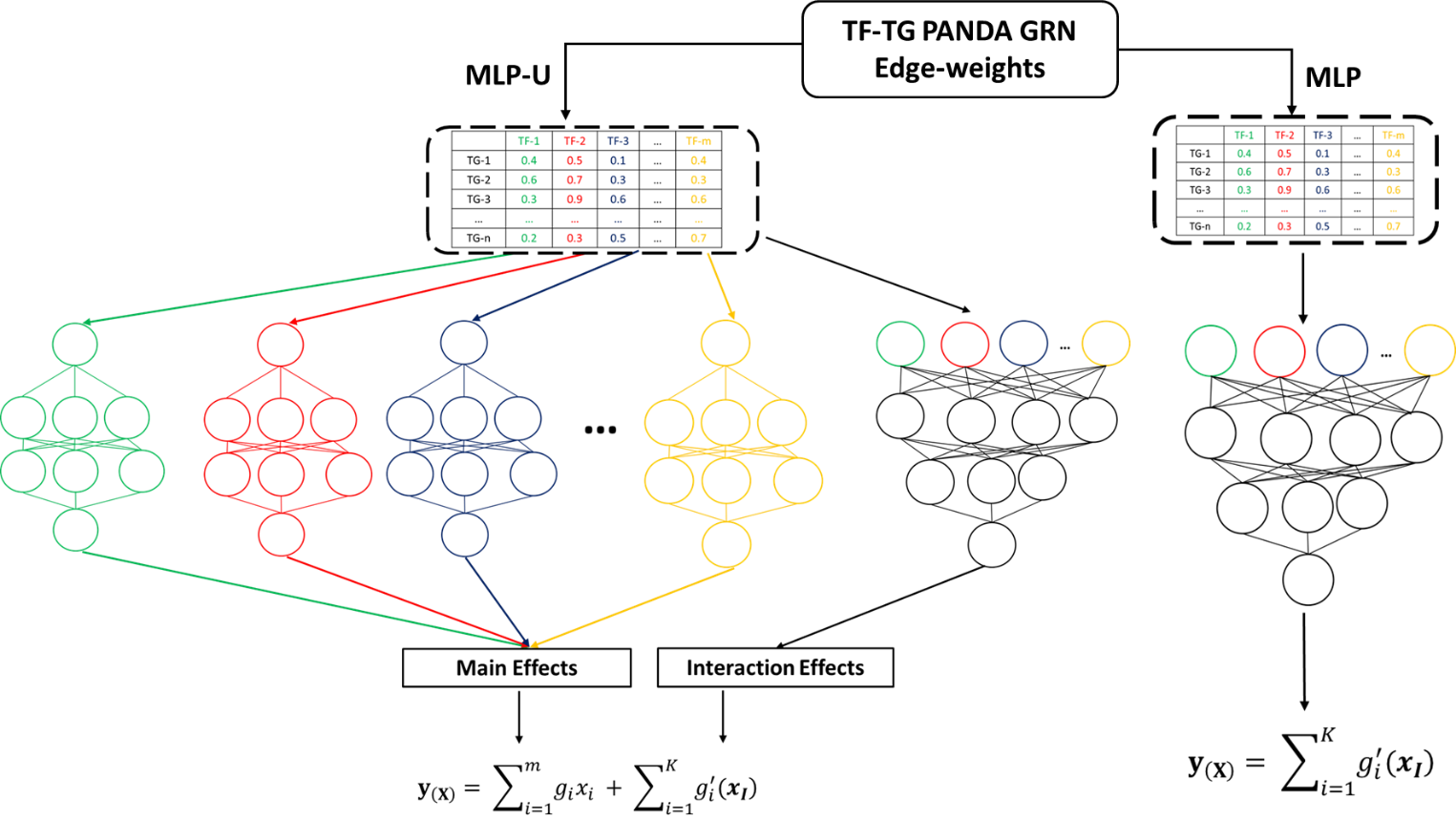

**Supplementary Figure S1:MLP-U and MLP architecture used in the paper. We trained two different types of MLP architectures in the paper: MLP-Uor MLP-Univariate(Left) and traditional MLP(Right). For the MLP-U models, we utilized individual TF edge-weights as inputs for the corresponding MLPs and trained them together with a traditional MLP receiving inputs corresponding to all the TFs. This ensemble model represented a generalized additive model where the main effects were derived from each individual MLP while the interaction effects were derived from traditional MLP. The traditional MLP model was trained without the individual MLPs and was assumed to just model the interaction effects. The modelling process involved partitioning the TG expression dataset into test and training sets using a 20-80 split and then training the models for 20 iterations. For the MLP-U model, we trained the individual univariate MLPs and the traditional MLP together via backpropagation.**

**Partitioning TG expression variance explained by the univariate and MLP components from the MLP-U models**

The MLP-U model architecture shown in ***Supplementary Figure S1*** contained individual univariate MLPs corresponding to each one of the 149 TFs receiving input features from these TFs as well as a traditional fully connected MLP receiving input features corresponding to all the TFs at the same time. Each MLP-U model was trained using the generalized additive equation shown in ***Supplementary Figure S1*** containing the univariate main effects as well as the interaction effects used to predict TG expression. In order to partition the variance in TG expression explained by these two components of the MLP-U models, we followed the following steps: 1) We obtained the learned MLP-U models trained from predicting TG expression for the 20 random states described in ***Supplementary table S2*** and extracted weights corresponding to the layers of the univariate MLPs as well as of the traditional MLP for each model. 2) We used the layer weights to generate two separate MLP models based on univariate and traditional MLP neural networks. 3) Lastly, we used these two models to independently predict expression for the test set TGs defined based on the random states provided in ***Supplementary table S2*** and for each prediction task we calculated the R^2^ that reflected the variance explained in the TG expression for the two models. We plotted these R^2^ for each prediction round for the two models in ***Supplementary Figure S2***

**Supplementary Figure S2: The boxplot shows the prediction performance calculated in the form of R^2^(variance explained) by predicting TG expression(N = 2,356) over 20 iterations using the univariate and the MLP components of the learned MLP-U models. (****p-value < 0.0001 calculated using paired t-test)**

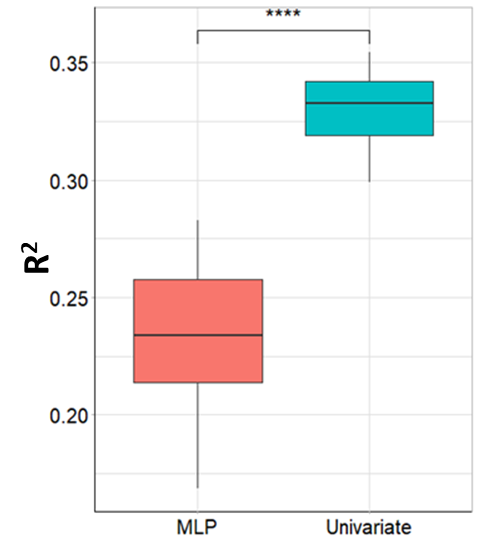

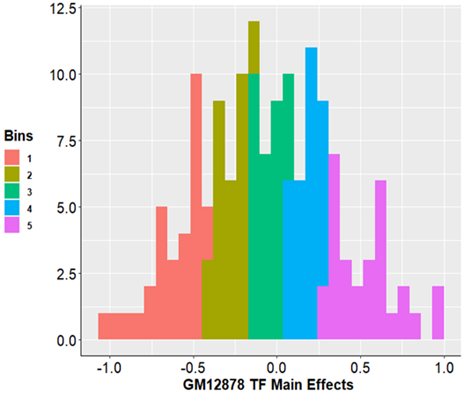

**Supplementary Figure S3: Histogram of the scaled main effects for each TF obtained after aggregating the layer weights of the MLP-U models corresponding to each TF across the 20 iterations. The histogram was further divided into 5 equal bins based on the scaled main effects.**

**Supplementary Figure S4: Barplot showing the nestedness for each higher-order(three-way or higher) TRM calculated as the proportion of all the possible pairwise interactions being also present in the detected set of TRMs. The nestedness of none of the higher order TRMs was 100%. In other words, none of the higher order TRMs could be completely explained by their subset pairwise interactions.**

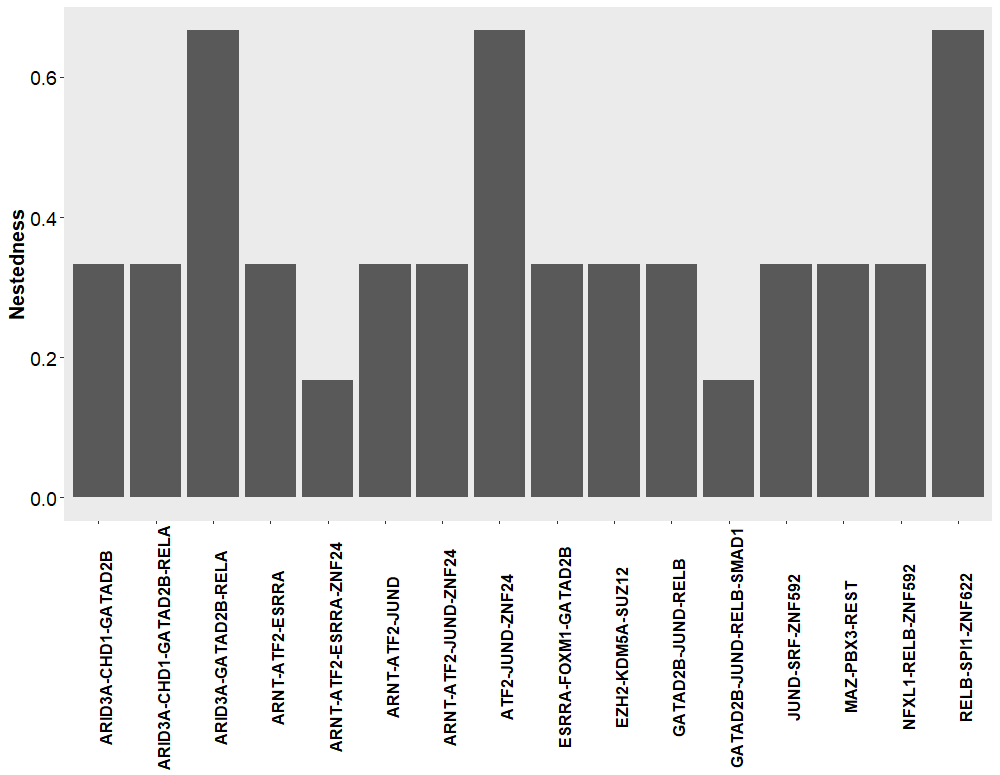

**Supplementary Figure S5: Barplot showing the mean log10 Hi-C contacts(1Kb resolution) between peak regions of the pairwise TRMs shaded according to the respective log2 NID scores across all the TG. We weren’t able to detect any HiC contacts between the peak pairs of the TRM SUZ12-ZNF274.**

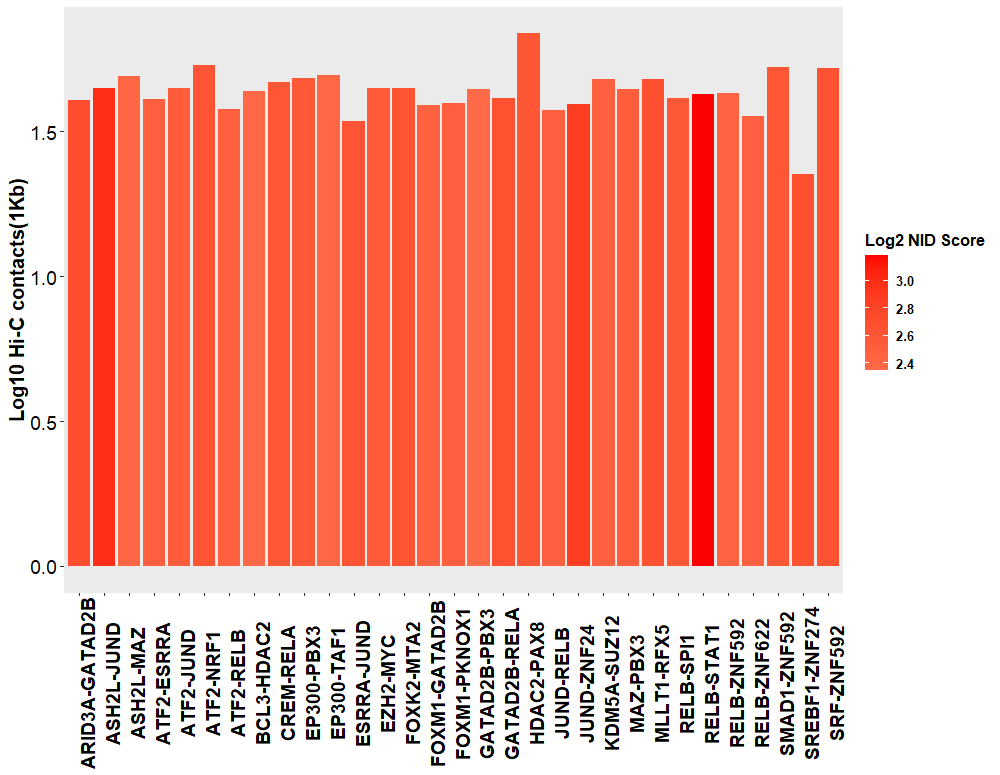

| **S1** | **ENCODE accessions for the GM12878 ChIP-Seq files used in the study** |
| --- | --- |
| **S2** | **PCC obtained from the prediction of the test set TGs during each of the 20 prediction runs with the random state used for each run for the three models.** |
| **S3A** | **Main effects for the 149 TFs derived from the learned MLP-U and ENET models along with the ranks based on these effects. We have also provided scaled version of the main effects and the difference in ranks(DIR) in this table.** |
| **S3B** | **NID and Log2NID scores for each one of the 48 TRMs identified in the paper** |
| **S4A** | **Two-by-two tables containing number of peak pairs with and without Hi-C contacts used for calculating p-value for Hi-C contact enrichment.** |
| **S4B** | **HiC contacts made by the TF peak pairs corresponding to the 31 pairwise TRMs present within the *cis*-regulatory region of TGs at 5Kb resolution.** |
| **S4C** | **Significantly co-localizing TF peak pairs corresponding to 6 pairwise TRMs within the *cis*-regulatory regions of TGs at the p-value threshold of 0.0001 identified using the SpaMo algorithm.** |
| **S5** | **Transcription regulatory programs used by each TRM for regulating the TGs along with the corresponding peak pairs.** |

***Descriptions of the supplementary tables attached with the manuscript***

**References**

[1]	C. A. Davis *et al.*, “The Encyclopedia of DNA elements (ENCODE): data portal update,” *Nucleic Acids Res.*, vol. 46, no. D1, pp. D794–D801, Nov. 2017, doi: 10.1093/nar/gkx1081.

[2]	S. Durinck *et al.*, “BioMart and Bioconductor: a powerful link between biological databases and microarray data analysis.,” *Bioinformatics*, vol. 21, no. 16, pp. 3439–3440, Aug. 2005, doi: 10.1093/bioinformatics/bti525.

[3]	O. Fornes *et al.*, “JASPAR 2020: update of the open-access database of transcription factor binding profiles,” *Nucleic Acids Res.*, vol. 48, no. D1, pp. D87–D92, Nov. 2019, doi: 10.1093/nar/gkz1001.

[4]	C. E. Grant, T. L. Bailey, and W. S. Noble, “FIMO: scanning for occurrences of a given motif,” *Bioinformatics*, vol. 27, no. 7, pp. 1017–1018, Feb. 2011, doi: 10.1093/bioinformatics/btr064.

[5]	T. Whitington, M. C. Frith, J. Johnson, and T. L. Bailey, “Inferring transcription factor complexes from ChIP-seq data,” *Nucleic Acids Res.*, vol. 39, no. 15, pp. e98–e98, May 2011, doi: 10.1093/nar/gkr341.

[6]	R. Oughtred *et al.*, “The BioGRID interaction database: 2019 update,” *Nucleic Acids Res.*, vol. 47, no. D1, pp. D529–D541, Nov. 2018, doi: 10.1093/nar/gky1079.

[7]	T. Lappalainen *et al.*, “Transcriptome and genome sequencing uncovers functional variation in humans,” *Nature*, vol. 501, no. 7468, pp. 506–511, 2013, doi: 10.1038/nature12531.

[8]	K. Glass, C. Huttenhower, J. Quackenbush, and G.-C. Yuan, “Passing Messages between Biological Networks to Refine Predicted Interactions,” *PLoS One*, vol. 8, no. 5, p. e64832, May 2013.

[9]	H. Zou and T. Hastie, “Regularization and Variable Selection via the Elastic Net,” *J. R. Stat. Soc. Ser. B (Statistical Methodol.*, vol. 67, no. 2, pp. 301–320, Jan. 2005.
